# nf_xpatial: A Reproducible Framework for Standardized Preprocessing and Clustering of Xenium Data

**DOI:** 10.64898/2026.08.25.747147

**Authors:** Luke A. Potter, Austyn Trull, Nilesh Kumar, Olivia R. Drake, Margareth Nogueira, Jamie Peters, Jasper A. Heinsbroek, Jeremy J. Day, Elizabeth A. Worthey, Lara Ianov

## Abstract

Recent advances in spatial transcriptomics have enabled the profiling of increasingly larger numbers of genes while retaining single-cell and subcellular resolution *in situ*. However, standardized bioinformatics workflows for analyzing these datasets have lagged behind, with existing pipelines focusing primarily on image processing and cell segmentation. To address this gap, we present nf_xpatial, a best-practices Nextflow pipeline for the downstream analysis of 10x Genomics Xenium data. The pipeline performs quality control, filtering, log and cell area normalization, multi-sample integration, and both expression-driven and spatially informed clustering across systematic parameter sweeps, allowing users to evaluate and compare clustering resolutions and spatial modeling parameters within a single reproducible run. Overall, nf_xpatial streamlines the processing of Xenium data from platform outputs to integrated single-cell and spatial clustering datasets, providing a standardized starting point from which biologists can finetune parameters and proceed to hypothesis-driven spatial analyses.

**Availability and implementation:** The source code and detailed documentation are freely available at https://github.com/U-BDS/nf_xpatial under the GPL-3 license.

**SUPPLEMENTARY INFORMATION:** Supplementary data is provided.

## INTRODUCTION

The application of spatial transcriptomics is rapidly advancing molecular biology, pathology, and systems biology by supporting the interrogation of gene expression within intact tissue architecture (Carstens, et al., 2024; Moses and Pachter, 2022; Wang, et al., 2025). Whereas earlier assays such as immunohistochemistry and RNA *in situ* hybridization were limited in target and multiplexing capacity, (Lowe, et al., 2017) sequencing-based platforms enable transcriptome-wide spatial profiling (Stahl, et al., 2016), and imaging-based platforms (e.g., MERSCOPE, CosMx SMI, and Xenium) quantify hundreds to thousands of targeted transcripts at single cell to subcellular resolution (Chen, et al., 2015; He, et al., 2022; Janesick, et al., 2023; Zhang, et al., 2021).

As these platforms expand in capabilities and generate increasingly large datasets, the need for standardized analytical workflows also grows. Vendor software (Bruker Spatial Biology, 2022; Janesick, et al., 2023) and community pipelines such as nf-core/sopa and nf-core/spatialaxe (Blampey, et al., 2024; Ewels, et al., 2020) primarily address image processing, transcript decoding, and cell segmentation, while robust and reproducible tertiary analysis remains underdeveloped. Existing approaches generally fall into three categories: analyses tailored to a specific study or biological question (Chan, et al., 2026; Vo, et al., 2025); pipelines that provide only partial tertiary analysis, often omitting spatial-specific steps (Bilous, et al., 2026; Blampey, et al., 2024; Ewels, et al., 2020); and proprietary end-to-end software with limited flexibility (10x Genomics, 2024; Bruker Spatial Biology, 2022; Janesick, et al., 2023). Notably, 10x Genomics’ software ecosystem illustrates this gap. While the Xenium Onboard Analysis performs primary data processing, and the Xenium Explorer provides visualization and exploratory analysis, neither offers reproducible, multi-sample tertiary analysis solutions required to derive biological insights. Consequently, users must perform substantial *ad hoc* analyses to effectively analyze Xenium data.

To address this gap, we present nf_xpatial, a modular and reproducible Nextflow workflow for Xenium data preprocessing. It standardizes quality control (including cell area and regionspecific filtering), applies log and cell area normalization, and supports multi-sample integration and systematic parameter exploration for both expression-driven cell-type and spatial-domain clustering within a single reproducible run.

## PIPELINE DESIGN AND IMPLEMENTATION

nf_xpatial is built with Nextflow DSL 2.0 (Di Tommaso, et al., 2017), with each process executed by R (R Core Team, 2025) within Docker and Singularity containers to ensure reproducibility and portability. It allows users to analyze the outputs from the Xenium platform, performing in-depth quality control, filtering, normalization, and clustering across systematic parameter sweeps (**Fig. 1**) with results consolidated into a report and compiled Seurat objects (Hao, et al., 2024). The following sections detail its components:

**Figure 1:**
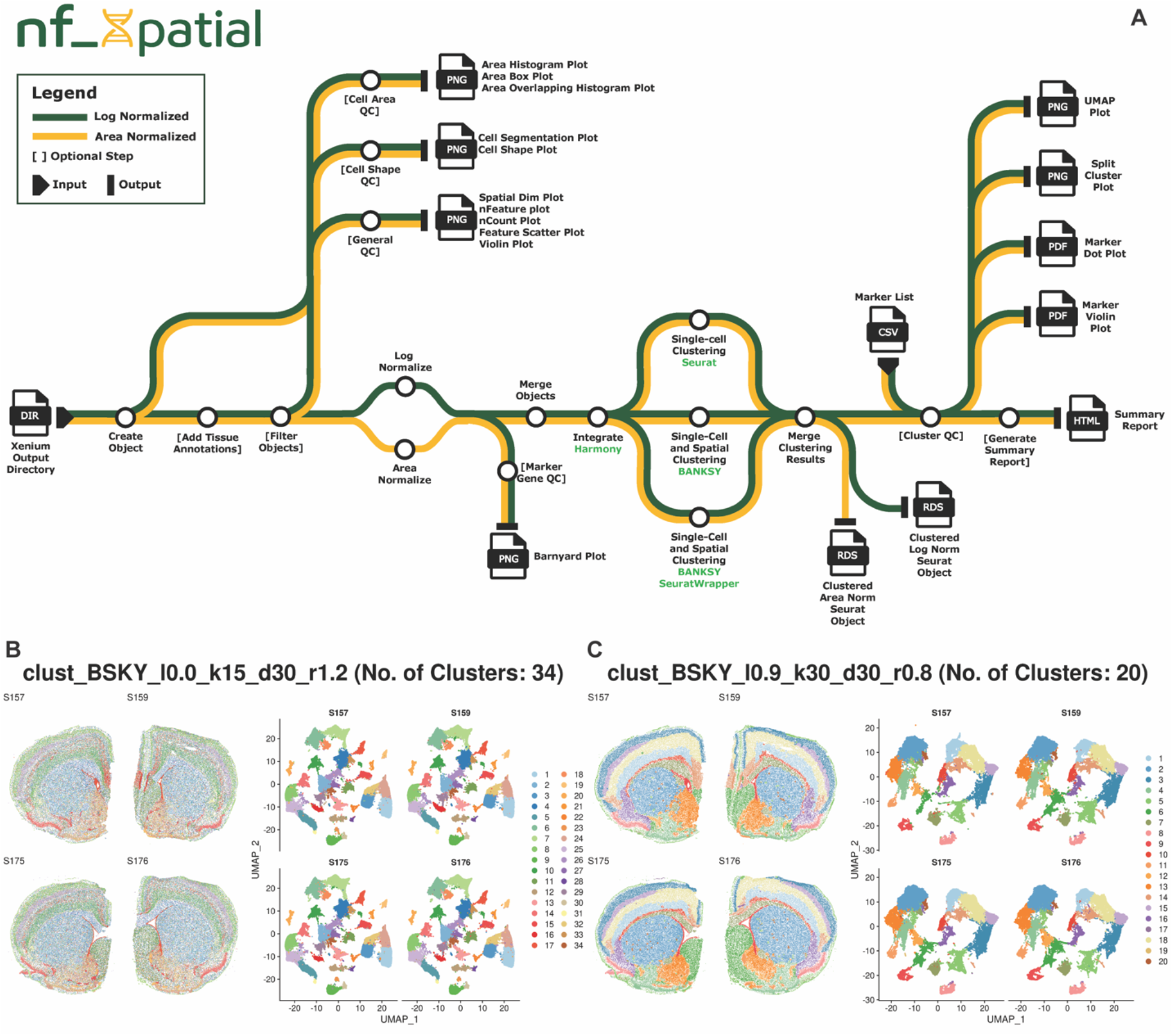
nf_xpatial workflow overview. nf_xpatial performs comprehensive preprocessing of Xenium spatial transcriptomics data. **A**. The diagram outlines the pathways to perform quality control, filtering, data normalization (area and log normalization), multi-sample integration, and systematic parameter exploration for both expression-driven cell type clustering and spatial domain clustering approaches. The deliverables include a comprehensive report and compiled Seurat objects. **B-C**. Representative outputs from the nf_xpatial parameter sweep when BANKSY is selected as the clustering method. Each panel title reflects the parameter combination used: BANKSY’s lambda (I), spatial neighbors (k), PCA dimensions (d), and resolution (r).

### Initial Processing

nf_xpatial accepts as input either the output directories created by the Xenium Onboard Analyzer or unprocessed per-sample Seurat objects, the latter accommodating upstream processing that a user may have performed, such as re-segmentation. Additionally, nf_xpatial accepts sample metadata which is added to the Seurat object(s) the pipeline delivers.

Optionally, the pipeline also accepts regions of interest (ROI) exported from the Xenium Explorer, where users can manually delineate and label ROIs within their tissue sections. If supplied, these ROIs are carried through the workflow and stored in the metadata of the resulting Seurat object. The same mechanism can be used to exclude ROIs, such as areas belonging to adjacent tissue or portions of the section that have folded onto themselves.

Following object creation and optional ROI removal, objects are filtered to remove low-quality cells using nFeature (detected features per cell), nCount (detected transcripts per cell), and spatial-specific thresholds such as cell area. All thresholds are exposed as pipeline parameters, providing granular control over the values applied and allowing individual filters to be skipped.

### Normalization

nf_xpatial supports two methods of normalization, which can be executed individually or in parallel. Log normalization is performed using Seurat, and scales gene expression counts for each cell based on total expression, multiplies the normalized values by a scaling factor, and applies a natural log transformation. Area normalization is implemented as a custom script based on the algorithm proposed by Atta et al. (2024). This method scales gene counts relative to the median cell area within each sample and applies a natural log-transformation on the result.

### Integration and Clustering

Following normalization, the data are merged and integrated using Harmony (Korsunsky, et al., 2019). Integrated data can then be clustered using one of two methods. Seurat clustering provides expression-driven cell type clustering, whereas BANKSY (Singhal, et al., 2024) supports both expression-based cell type clustering and spatial domain clustering. The pipeline provides two ways to invoke BANKSY: 1. by using BANKSY through the Seurat Wrapper or 2. by using the base BANKSY R package. All clustering methods accept a comma-delimited list of parameter values (e.g., dimension, resolution, and for BANKSY, lambda (*λ*) and k_geom) to allow for the evaluation of different cluster configurations. Upon completion, the clustering results generated for each normalization are merged into a single object.

### Quality Control, Visualization, and Report

Quality control (QC) figures are generated throughout the workflow. Spatial dimension plots, per-cell feature (nFeature) and transcript (nCount) distributions, feature scatter plots, and violin plots are produced for both raw and filtered data, enabling data quality to be evaluated prior to normalization and clustering.

Additional spatially specific QC figures are generated to facilitate the evaluation of cell segmentation quality and detection of tissue dissimilarities. When multimodal segmentation is applied, cell segmentation metrics are summarized into a single plot, allowing segmentation quality assessment across all samples in a run. Cell shape plots similarly summarize the estimated cell shape for each sample into a single plot, classifying cells as “circular,” “polygonal,” or “elongated” based on roundness and aspect ratio. Cell area plots compare cell area distributions across samples, with cells flagged as outliers using an interquartile range (IQR) criterion: those with areas below Q1 - 1.5×IQR or above Q3 + 1.5×IQR, where Q1 and Q3 are the 25^th^ and 75^th^ percentiles of cell area by default. Flagged cells are reported for inspection rather than automatically removed, allowing users to determine the most appropriate cutoffs. Additionally, barnyard plots compare the expression of gene pairs across all cells within a tissue sample and provide an additional QC metric.

Finally, UMAP visualizations and marker gene dot or violin plots are generated to evaluate clustering results and guide parameter selection for downstream analyses. These outputs, together with selected figures from above, are compiled into a single summary report.

## APPLICATION

To demonstrate the utility of nf_xpatial, we applied the workflow to an in-house Xenium mouse brain dataset comprising four coronal hemisections (one per animal) assayed using a 347-gene probe set (10x Genomics Mouse Brain Panel and 100 custom genes). To benchmark computational resource requirements for Xenium Prime 5K data, we also ran nf_xpatial on six FFPE non-small-cell lung cancer samples from Bilous et al (2026) (see Supplemental Information).

Applying nf_xpatial to the mouse brain dataset demonstrates the workflow’s comprehensive QC capabilities, including preliminary metrics combining standard single cell metrics (transcript and feature distributions, their correlation, and their spatial distribution across the tissue), with imaging-specific assessment of cell area QC and per cell area distribution (**Supplemental Fig. 1**). When multimodal cell segmentation is applied, nf_xpatial also generates cohort-level summaries to aid in rapid assessment of the distribution of segmentation methods across samples (**Supplemental Fig. 2A**). Similarly, nf_xpatial classifies cell shapes into three categories (circular, elongated, and polygonal) to aid in assessing cohort similarity for QC purposes (**Supplemental Fig. 2B**). In either metric, large deviations between samples may be indicative of QC issues. The workflow additionally supports visualization of paired marker co-expression, providing an orthogonal check for transcript misassignment between neighboring cells for any markers that may be mutually exclusive (**Supplemental Fig. 2C-D**).

Here, we enabled both log and cell area normalization and all three clustering workflows (Seurat, BANKSY, and the BANKSY Seurat wrapper) across a sweep of parameters including dimension, clustering resolution, and, for BANKSY, the spatial weight *λ*, and the neighborhood size, k_geom. Although all three approaches are shown here for the mouse brain dataset, users may enable one individual method or any combination of the three. Results are summarized as integrated UMAPs, split-cluster plots showing per-sample UMAPs and spatial dimension plots, and marker expression dot plots (**Fig. 1B-C** and **Supplemental Fig. 3A-C**).

Although nf_xpatial does not perform automated cell-type annotation, its output objects are compatible with downstream annotation workflows. To determine whether the resulting clustering recovers biologically coherent populations, we annotated one expression-driven result using canonical markers (which nf_xpatial visualizes for each parameter set) and labels generated by MapMyCells (Daniel, et al., 2026). This analysis recovered the expected neuronal and nonneuronal populations, including cortical layer-specific glutamatergic neurons, striatal medium spiny neurons, GABAergic interneuron subtypes, and glial cells (**Supplemental Fig. 3D-E and 4**).

Lastly, because runtime and resource requirements scale with cell number, sample count, and the number of parameter sets evaluated, we profiled computational requirements. **Supplementary Figs. 5** and **6** summarize the peak memory usage and maximum wall-clock runtime of each process for the mouse brain and Bilous et al. 5K datasets, respectively (see Supplemental Extended Benchmark).

## CONCLUSION

Here we introduce nf_xpatial, a standardized and reproducible framework for processing 10x Genomics Xenium data. Our pipeline supports in-depth QC, filtering, log and cell area normalization, sample integration, and both expression-driven cell type and spatial domain clustering across systematic parameter sweeps. Results are consolidated into a summary report and analysis-ready Seurat objects. By automating these steps, nf_xpatial reduces the manual effort and *ad-hoc* code typically required for Xenium analysis, enabling researchers to progress to hypothesis testing more quickly, while retaining granular control through its modular and configurable design.

Planned enhancements include the addition of more computationally efficient clustering algorithms. For instance, the BANKSY developers are evaluating lazy sparse matrix multiplication for PCA, which avoids construction of the full BANKSY matrix and should reduce runtime and memory use for large datasets. We are evaluating this approach within nf_xpptial and will incorporate it when it becomes available. We are also exploring cell-label transfer to streamline annotation. These improvements will become increasingly important as whole-transcriptome *in situ* platforms, such as the 10x Genomics Atera, generate increasingly larger and more complex datasets. Importantly, the modular design of nf_xpatial ideally positions it to adapt to these emerging technologies.

Overall, nf_xpatial provides a robust, standardized analysis solution from raw Xenium output to results ready for cell type annotation and spatial domain identification, allowing users to focus on biological interpretation.

## AUTHOR CONTRIBUTIONS

J.P., J.A.H., J.J.D., L.I., E.A.W. obtained funding. L A P., A.T., N.K., L.I. designed and developed nf_xpatial. O.R.D., M.N., J.A.H performed sample collection and preparation of the mouse brain dataset. LAP., A.T., L.I. performed validation, benchmarking, and data visualization as detailed. L A P, A.T., L.I. drafted the manuscript. L.I. supervised all work. N.K., O.R.D., M.N., J.P., J.A.H., J.J.D., E.A.W. provided feedback and revised the manuscript. All authors approved the final manuscript.

## ACKNOWLEDGMENTS

We acknowledge support from the University of Alabama at Birmingham Biological Data Science Core, RRID:SCR_021766. We also acknowledge support from the nf-core community for developing and maintaining the nf-core template, which was implemented in nf_xpatial. The authors gratefully acknowledge the resources provided by the University of Alabama at Birmingham IT-Research Computing for high-performance computing (HPC) support and CPU time on the Cheaha compute cluster.

## SUPPLEMENTAL INFORMATION

### EXTENDED RESULTS

#### Supplemental figures

**Supplementary Figure 1:**
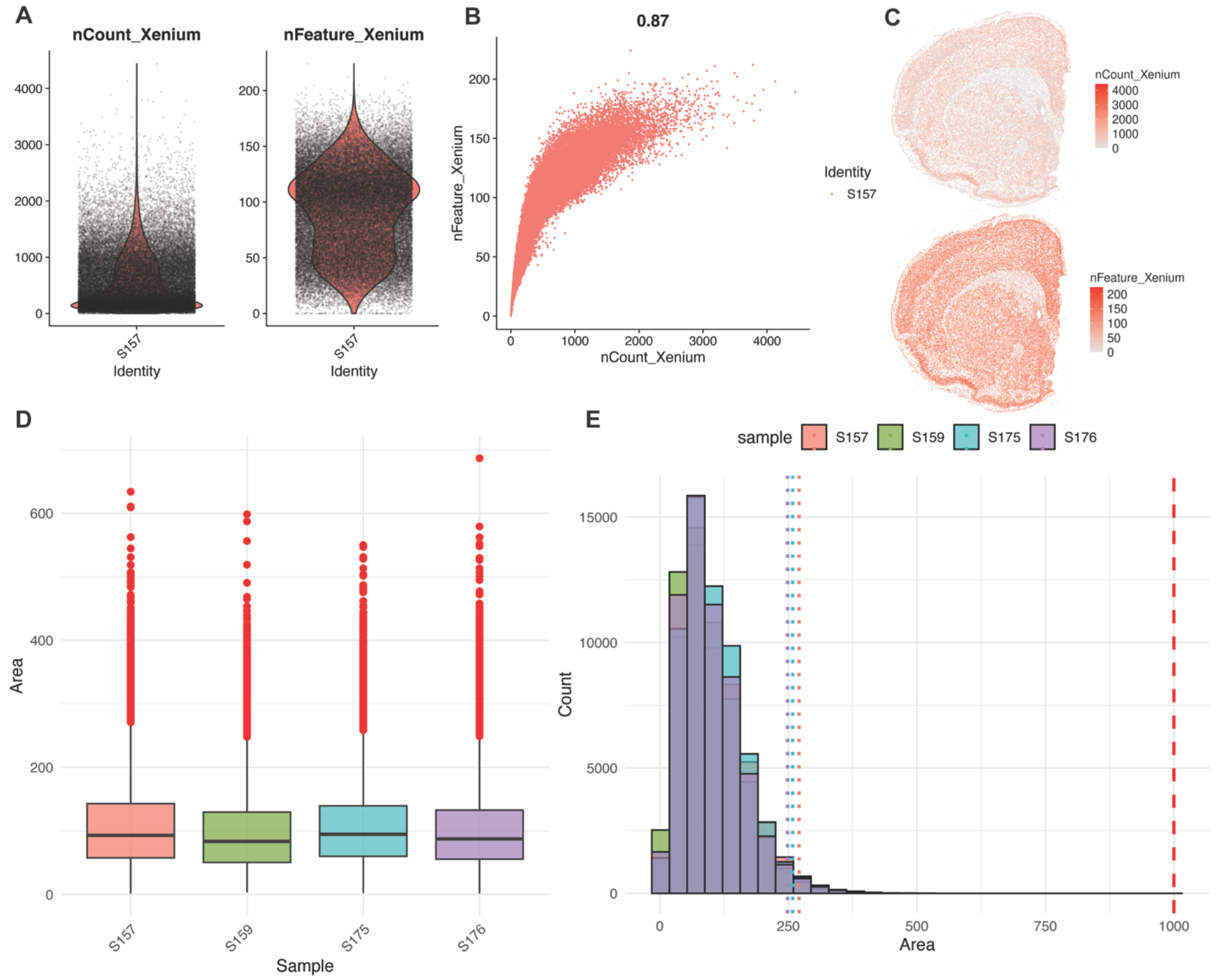
Single-cell QCs and cell area QCs. **A-C:** Representative outputs (of a single sample) of standard single-cell metrics including **A**. Violin plots of transcript counts per cell (nCount_Xenium) and detected features per cell (nFeatures_Xenium), with individual cells overlaid as points **B**. Scatter plot of transcript counts and features (Peason correlation shown above panel) and **C**. Spatial feature plot of the distribution of transcript counts and features across the tissue section. **D-E**. Representative outputs of the cell area distribution across all samples including **D**. Box plots of the cell area per sample (red points indicate cells which have been flagged as outliers defined as cell area below Q1 - 1.5xIQR or above Q3 + 1.5xIQR) **E**. Histograms of the distributions of the cell areas across all samples overlaid. The vertical dashed red line denotes a user-specific threshhold while the dotted lines colored by sample denote each sample’s upper outlier threshhold.

**Supplementary Figure 2:**
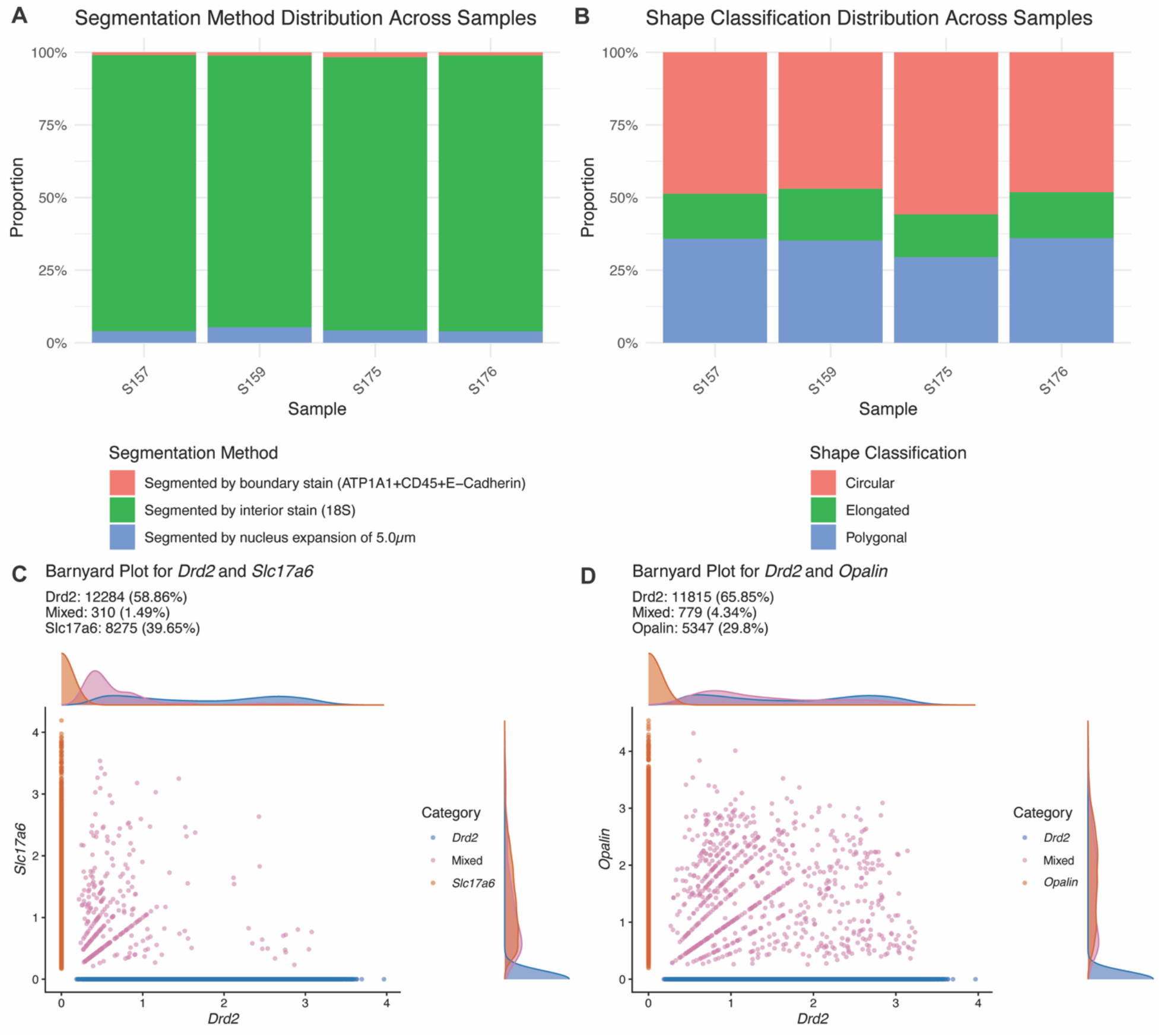
Aggregated cross-sample metrics for rapid quality assessment and segmentation quality. **A-B**. Representative QC outputs summarized across all samples, enabling deviating samples to be identified at a glance. **A**. Proportion of cells resolved by each Xenium multimodal segmentation method shown per sample. **B**. Proportion of cells assigned to each shape classification (circular, elongated, polygonal). In both panels, comparable tissue sections processed under equivalent conditions are expected 10 yield similar proportions across samples. A sample whose composition departs substantially from the others should be flagged as a potential QC failure. **C-D**. Barnyard plots of per-cell co-expression of *Drd2 /Sld7a6* (0) and *Drd2 /Opalin* (**D**). Cells falling along either axis show the expected mutually exclusive expression, whereas double-positive cells (purple) indicate transcript misassignment between adjacent cells. A low proportion of double-positive is expected for accurate segmentation.

**Supplementary Figure 3:**
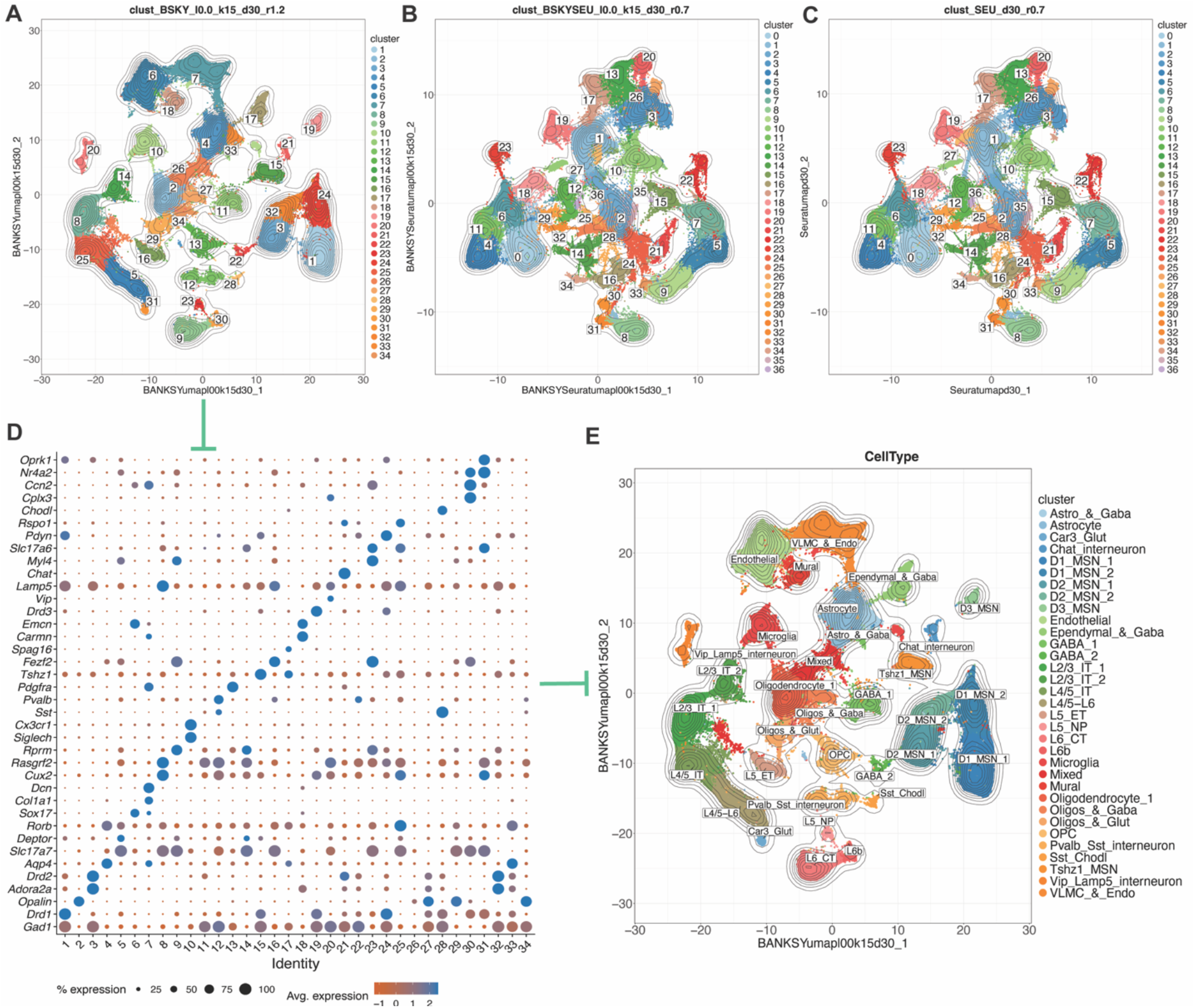
Overview of the three clustering approaches implemented in nf_xpatial and downstream cell type annotation. Representative outputs of the expression-driven cell type clustering from the mouse brain dataset. **A-C**. UMAP embeddings of cells clustered using (A) BANKSY, (B) the BANKSY Seurat wrapper, and (C) Seurat, each shown at a representative parameter combination (indicated in the panel titles). Cells are colored by cluster assignment, with cluster labels overlaid; contour lines indicate cell density. **D**. Dot plot of selected marker genes across the clusters obtained from the BANKSY workflow in (A). **E**. UMAP from (A), with clusters annotated to cell type identities on the basis of the marker expression shown in (D) and MapMyCells label transfer (Supplemental Figure 4). Annotations recover expected neuronal and non-neuronal populations, including cortical layer-specific glutamatergic neurons, striatal medium spiny neurons, GABAergic interneuron subtypes, and glial populations.

**Supplementary Figure 4:**
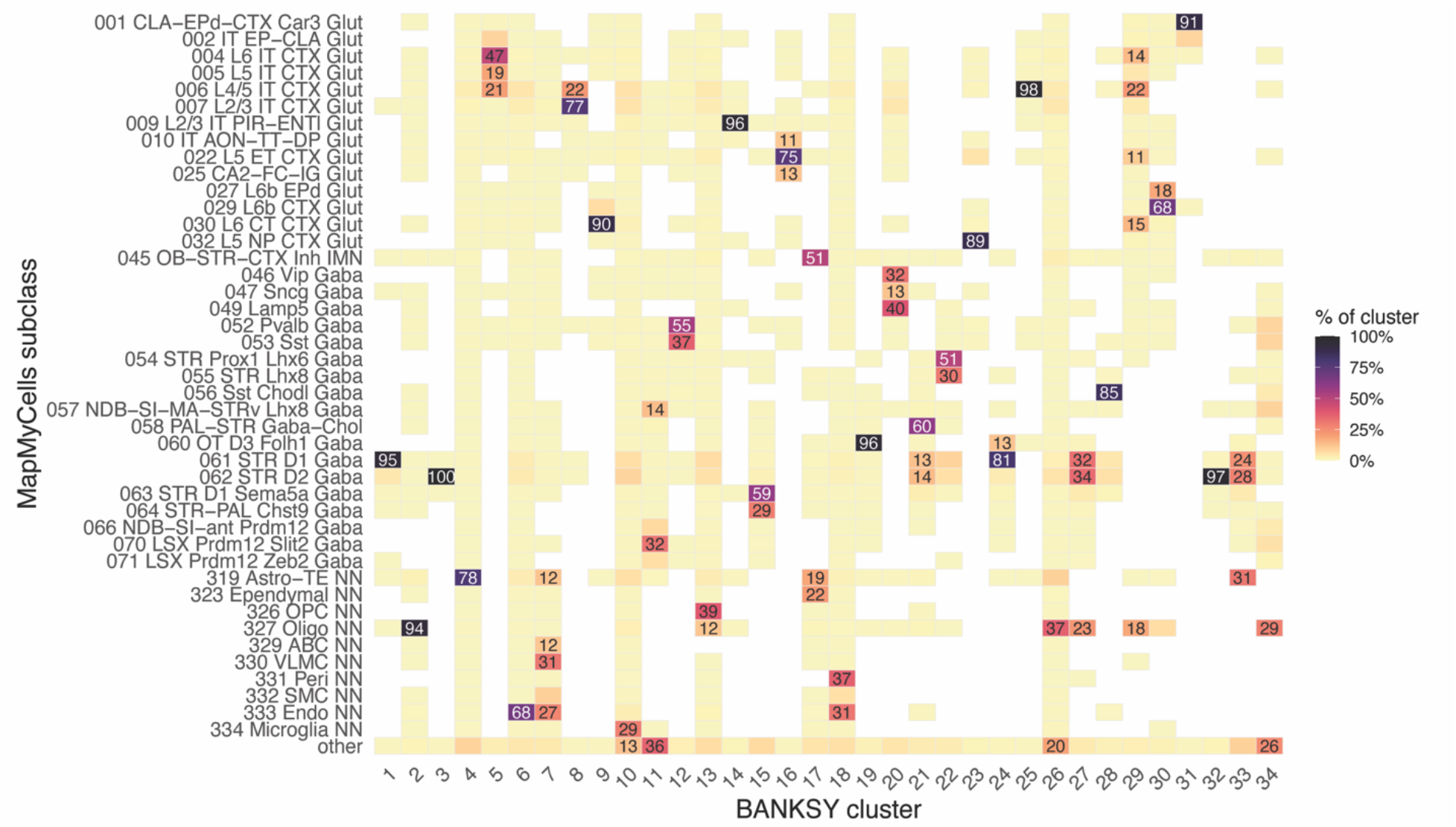
MapMyCells subclass composition. Heatmap showing, for each unsupervised BANKSY cluster (Supplemental 3A), the proportion of the cells assigned to each whole mouse brain taxonomy subclass by MapMyCells. Tile fill encodes the percentage of the cluster’s cells mapping 10 that subclass (color bar: light =low dark - high), and values ≥ 10% are printed. Subclasses reaching ≥ 10% at least one cluster are shown individually; all others are aggregated as “other.”

## EXTENDED BENCHMARK

nf_xpatial was benchmarked across the mouse brain and the Bilous *et al* FFPE non-small-cell lung cancer (NSCLC) 5K datasets to quantify per-process runtime and peak memory, as well as total runtime across varying parameter sweep configurations and data size. Total runtime scales with both the number of parameter combinations assessed, the number of clustering strategies enabled, and the number of normalization strategies enabled. The benchmarks presented here therefore serve as a practical guide when considering how broad a parameter sweep to run for a given dataset size, which clustering method to choose, and whether to enable one or both normalization strategies.

In the mouse brain dataset, nf_xpatial was executed across the following parameter sets: Seurat clustering at dimensions of 25 and 30 and resolutions spanning 0.3-0.8 (12 combinations); and both BANKSY and the BANKSY Seurat wrapper at a dimension of 30, resolutions spanning 0.4-1.2, k_geom of 15 and 30, and *λ* of 0.0, 0.8 and 0.9, yielding 54 combinations each. Each combination was in turn run under both normalization strategies, totaling 240 parameter runs (120 per normalization: 12 Seurat + 54 BANKSY + 54 BANKSY Seurat wrapper). The total runtime for this run was 4h 43m, and the per-process maximum runtime and peak memory are presented in **Supplementary Figure 5**. Likewise, in the Bilous *et al* NSCLC 5K dataset, nf_xpatial was executed with the following parameters: Seurat clustering at dimensions of 25 and 30 and resolutions spanning 0.4-0.7 (8 combinations); and both BANKSY and the BANKSY Seurat wrapper at a dimension of 20 and 30, resolutions spanning 0.5-1.5, k_geom of 30, and *λ* of 0.0, 0.8 and 0.9, yielding 66 combinations each. Each combination was in turn run under both normalization strategies, totaling 280 parameter runs (140 per normalization: 8 Seurat + 66 BANKSY + 66 BANKSY Seurat wrapper). The total runtime for this run was 16h 55m, and the per-process maximum runtime and peak memory are shown in **Supplementary Figure 6**. Notably, in the NSCLC 5K dataset, BANKSY required longer runtimes than the BANKSY Seurat wrapper, suggesting that the latter may be preferable under constrained computational resources, or when a broad parameter sweep is desired.

**Supplementary Figure 5:**
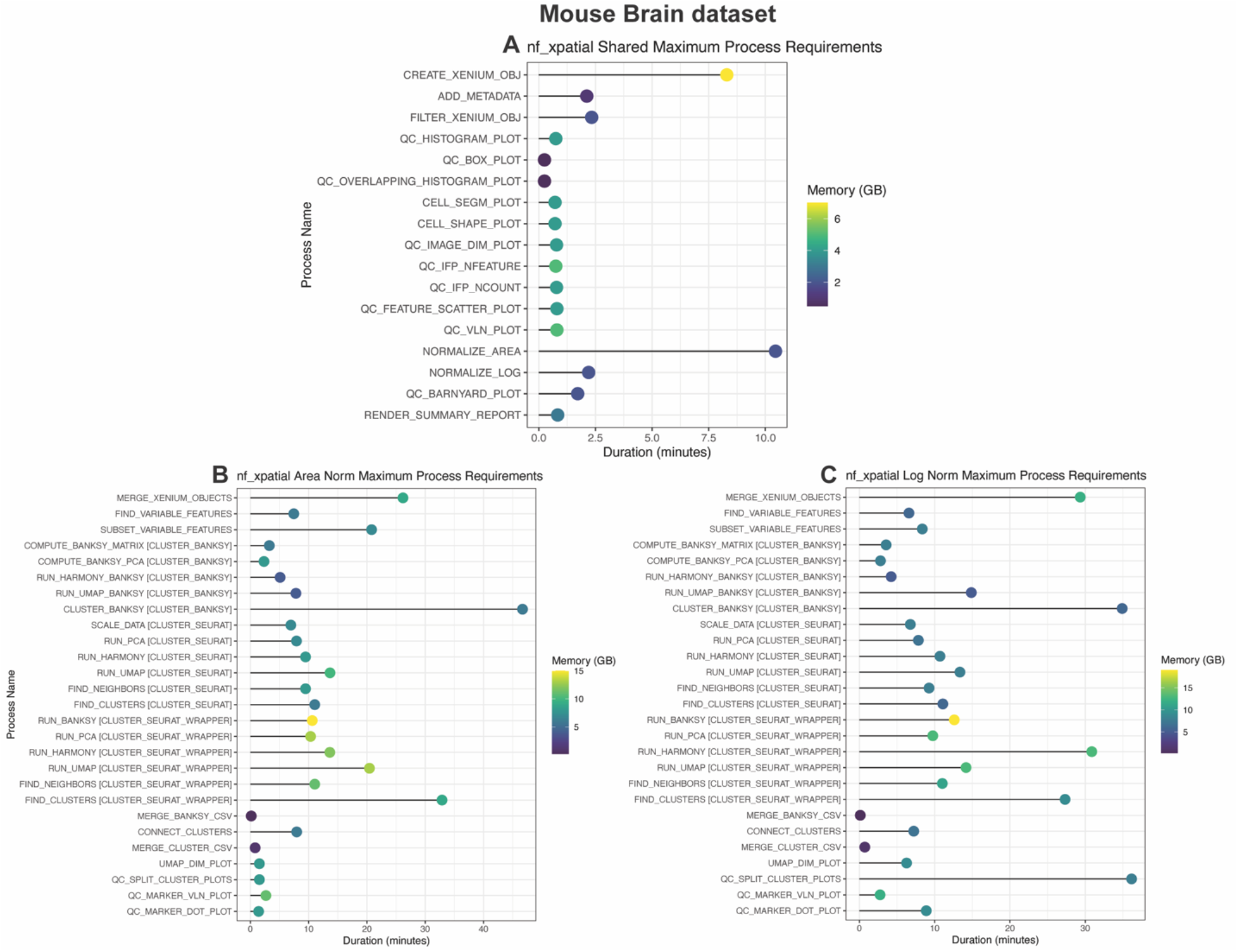
nf_xpatial benchmarks for the Mouse Brain dataset. rime (in minutes) and memory (in GB) requirements for processes shared across normalization methods (A) or normalization-specific processes (B: area normalization and C: log normalization).

**Supplementary Figure 6:**
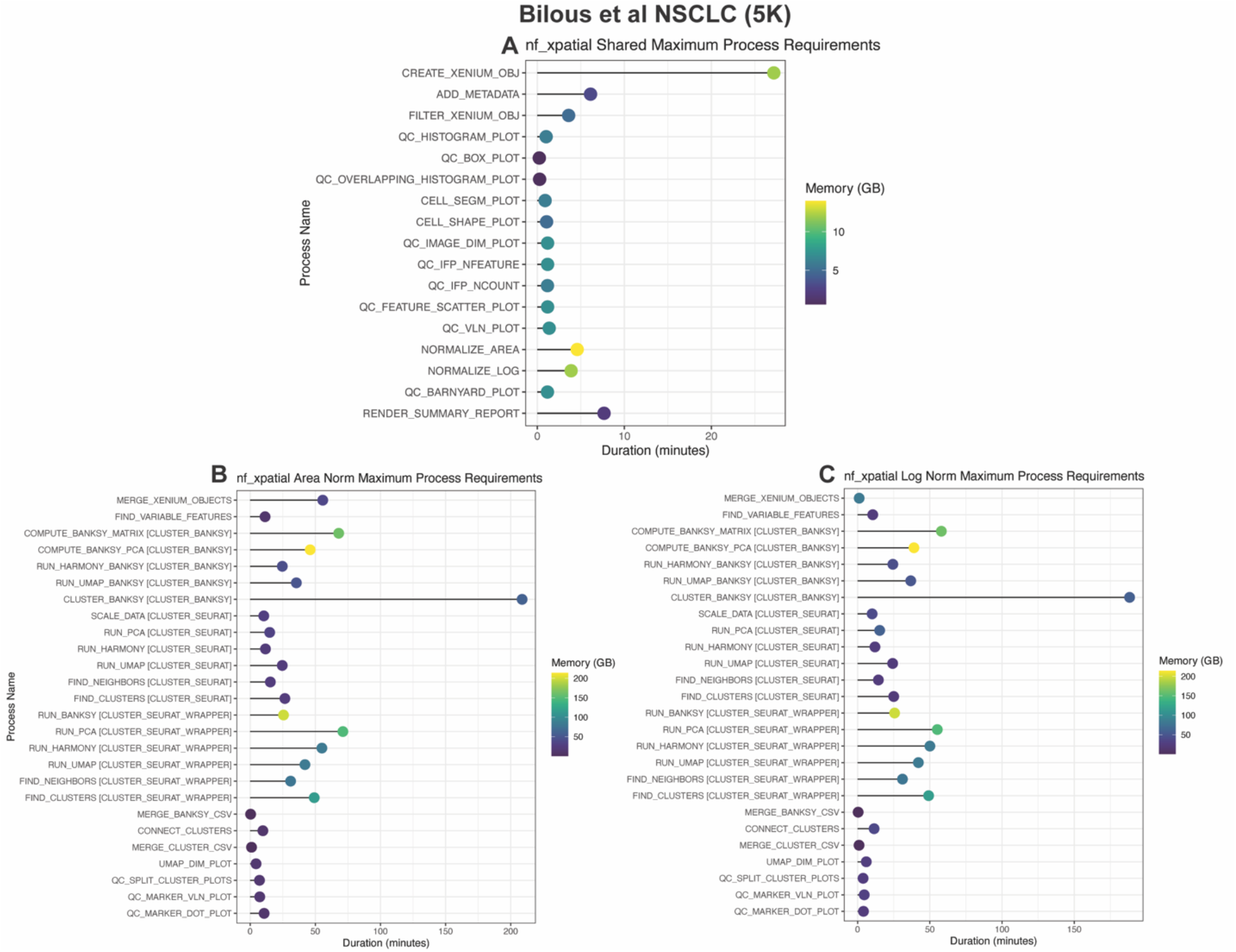
nf_xpatial benchmarks for the Bilous et al. NSCLC dataset. Time (in minutes) and memory (in GB) requirements for processes shared across normalization methods (A) or normalization-specific processes (B: area normalization and c: log normalization).

## METHODS

### Sample collection (mouse brain dataset)

One male and three female mice were euthanized by rapid decapitation, and whole brains were rapidly extracted. Brains were briefly rinsed in ice cold sterile PBS on ice, hemisected with a sterile razor blade (Fisher Scientific, 17-989-010), and immediately flash frozen in 2-methyl butane on dry ice. Brains were kept at −80°C until further processing. One hemisphere from each animal was sectioned at 10μm using a Leica CM 1860 cryostat (Leica Biosystems, Deer Park, IL, USA). One coronal hemisection per animal corresponding to approximately Bregma 11.10 mm was mounted onto a Xenium slide (10x Genomics, PN-1000460). Slides were stored at −80°C for no longer than 24 hours before further processing with the Xenium workflow.

### *In situ* spatial transcriptomic profiling

Xenium sample processing was performed by the Single Cell and Flow Cytometry Core at UAB. Spatial transcriptomic profiling was conducted using the Xenium Mouse Gene Expression Panel (10x Genomics, PN-1000462) supplemented with a custom gene panel (Supplemental Table 1) for a total of 347 probes. Tissue sections underwent fixation and permeabilization (protocol CG000579 Rev E) followed by probe hybridization, ligation, and amplification (protocol CG000749 Rev B). The slides were then placed in the Xenium analyzer for cycles of fluorescent probe hybridization, imaging, and fluorophore removal (protocol CG000584 Rev G). Fluorescent signals were registered across four imaging channels to generate spatially resolved transcript counts and positional barcode matrices for downstream analysis. Cell segmentation was performed using Xenium’s multimodal segmentation pipeline, incorporating membrane boundary staining (ATP1A1, CD45, E-Cadherin), interior stain (18S rRNA), and/or nucleus expansion of 5.0μm.

### Public Dataset

The Xenium Prime (5K gene panel) samples are derived from Bilous et al., and they include six non-small-cell lung cancer samples (available at the following GEO accession number: GSE311609). The sample numbers corresponding to the six Xenium Prime samples are: GSM9509134, GSM9509135, GSM9509136, GSM9509137, GSM9509138, and GSM9509139.

### nf_xpatial

nf_xpatial version 1.0.0 was used to analyze all datasets. Further, both normalization methods as well as all three clustering methods were enabled for benchmarking purposes. The parameter sets for each respective dataset are outlined under “Extended Benchmark”. Cells which contained fewer than 10 nCounts or 10 nFeatures were removed. Lastly, for the mouse brain dataset, *Mbp* was removed (via nf_xpatial’s flag ‘filter features’) due to molecular overcrowding. The scripts, parameters, and custom configuration files for each dataset can be found within the following repository: https://github.com/U-BDS/nf_xpatialjraper

### MapMyCells

For the mouse brain dataset, in addition to marker expression, which is provided by nf_xpatial, cell label transfer was performed using MapMyCells under the following configurations: Reference Taxonomy: 10x Whole Mouse Brain (CCN20230722) and Mapping Algorithm: Hierarchical Mapping. The results were imported into the nf_xpatial Seurat objects and summarized per cluster at the subclass level (**Supplementary Fig. 4**). Subclasses accounting for ≥10% of cells within a cluster are shown individually in the heatmap; all remaining subclasses are aggregated as “other.”

## AVAILABILITY OF DATA AND MATERIALS

nf_xpatial, including documentation, is available at https://github.com/U-BDS/nf_xpatial under the GPL-3 license. All dataset sources have been disclosed under the methods section. The mouse brain data has been deposited in GEO under the following accession number: GSE342202

